# Protection induced by anti-PD-1 and anti-PD-L1 treatment in *Leishmania amazonensis*-infected BALB/c mice

**DOI:** 10.1101/721894

**Authors:** Alessandra M. da Fonseca-Martins, Tadeu D. Ramos, Juliana E.S. Pratti, Luan Firmino-Cruz, Daniel Claudio Oliveira Gomes, Lynn Soong, Elvira M. Saraiva, Herbert L. de Matos Guedes

## Abstract

Leishmaniasis is a neglected disease, for which current treatment presents numerous issues. *Leishmania amazonensis* is the etiological agent of cutaneous and diffuse cutaneous leishmaniasis. The roles of the programmed death-1 (PD-1) receptor on lymphocytes and its ligand (PD-L1) on antigen-presenting cells have been well studied in tumor and other infection models; but little is known about their roles in non-healing cutaneous leishmaniasis. Our previous report of *L. amazonensis*-induced PD-L1 expression on dendritic cells, in combination with decreased IFN-γ production by CD4^+^ T cells in C57BL/6 mice, led to a hypothesis that the formation of the PD-1/PD-L1 complex contributes to down-modulation of immune responses, especially T cell suppression, enabling parasite survival and persistence. In this study, we tested the therapeutic potential of anti-PD-1 and anti-PD-L1 monoclonal antibodies (MoAbs) against a non-healing *L. amazonensis* infection in BALB/c mice. We observed that *L. amazonensis* induced PD-1 expression on both CD4^+^ and CD8^+^ T cells, and that anti-PD-1 and anti-PD-L1 treatment significantly increased IFN-γ-producing CD4^+^ and CD8^+^ T cells, respectively. Compared with infection controls, mice that received treatment with anti-PD-1 and anti-PD-L1, but not anti-PD-L2, displayed bigger lesions with significantly lower parasite loads. Treatment did not affect anti-*Leishmania* antibody or IL-10 production, but anti-PD-1 treatment reduced both IL-4 and TGF-β production. Together, our results highlight the therapeutic potential of an anti-PD-1-based treatment in promoting the reinvigoration of T cells for the control of parasite burden.

## INTRODUCTION

Leishmaniasis is a disease with global public health concerns, particularly in poor communities. Conventional treatments arose in the 1940s; however, many problems are associated with these medications, such as high toxicity, adverse side effects, and the increased incidence of drug-resistant parasites. At present, there are no vaccines available for human use, which makes the search for new molecular targets highly necessary^1^. *Leishmania amazonensis* infection can cause a diverse spectrum of the disease, including cutaneous (the most common), mucosal, and visceral leishmaniasis, as well as diffuse cutaneous leishmaniasis that is refractory to the conventional treatment^2^.

The programmed death-ligand 1 (PD-L1), a cell surface glycoprotein belonging to the B7 family is expressed on antigen-presenting cells such as neutrophils, macrophages, and dendritic cells. PD-L1 binds to the PD-1 receptor, which belongs to the CD28 family and is expressed on T cells, B cells, and myeloid cells^3-5^. The PD-1/PD-L1 interaction leads to the suppression of T cells by affecting the gradual loss of cell activities including cytokine secretion (IFN-γ, IL-2, TNF-α), decreasing the proliferative capacity, and finally, inducing T cell apoptosis.^6,7^ The PD-L1 receptor is widely discussed in oncological studies, as it is selectively expressed in many tumors^4,8,9^ and in cells within the tumor microenvironment in response to inflammatory stimuli.^10^ PD-L1 is positively regulated in solid tumors, where it can inhibit cytokine production and the cytolytic activity of PD-1-expressing CD4^+^ and CD8^+^ T cells.^4,11,12^ PD-1/PD-L1-based monoclonal antibody (MoAb) therapy is currently in phase III clinical trials with promising results for treatment against bladder carcinoma^13^ and lung cancer^14^. Programmed death-ligand 2 (PD-L2) is also a cell surface glycoprotein in the B7 family and plays a role similar to PD-L1, because it inhibits T cell function by binding PD-1 to the controversy in different models. T cell suppression is also reversed when the receptor is blocked by a specific antibody, for example, in inducing oral tolerance.^15-17^

It has been shown that PD-1/PD-L1-mediated cellular exhaustion also occurs during the progression of chronic infectious diseases caused by viruses or protozoan parasites, such as AIDS, toxoplasmosis, and cutaneous leishmaniasis. ^15,18-20^ Liang and colleagues have reported that *L. mexicana*-infected PD-L1^−/−^ mice have increased production of IFN-γ in T cells, reduced disease progression, and greater control of the parasite load when compared with infected wild-type (WT) mice. In PD-L2^−/−^ mice, however, there was an increased lesion size and increased parasite load compared to WT mice, which implies there are differing roles for PD-L1 and PD-L2 in regulating IFN-γ production. These results also suggest the participation only of PD-L1 in the T exhaustion process during *L. mexicana* infection.^15^

We have demonstrated that *L. amazonensis* infection of bone marrow-derived dendritic cells (BMDCs) from C57BL/6 and BALB/c mice induced PD-L1 expression. Furthermore, *L. amazonensis* infection of C57BL/6 mice induced suppression of the immune response through impaired IFN-γ-producing CD4^+^ T cells, which was not observed in PD-L1^−/−^ mice.^21^ In C57BL/6 mice, the lesion reaches a peak of growth during *L. amazonensis* infection, followed by a process of resolution of the disease, which shows that this mouse strain has a higher resistance to the parasite. In contrast, BALB/c mice develop a progressive lesion without healing, suggesting a greater susceptibility of these animals. Thus, we hypothesize that the use of anti-PD-1 and anti-PD-L1 MoAbs would have the potential to reverse the T cell suppression phenotype observed in BALB/c mice. Therefore, here we investigate the expression of PD-1 and PD-L1 upon *L. amazonensis* infection in BALB/c mice, and evaluate the use of MoAbs against PD-1, PD-L1 and PD-L2 as therapies for the severe form of leishmaniasis caused by *L. amazonensis*.

## MATERIALS AND METHODS

### Experimental Animals

Female BALB/c mice, 6-8 weeks old, from the Núcleo de Animais de Laboratório (Universidade Federal Fluminense, Rio de Janeiro, Brazil), were housed in Ventilife mini-isolators (Alesco, Brazil) and kept under controlled temperature and light conditions. All of the animal experiments were performed in strict accordance with the Brazilian animal protection law (Lei Arouca number 11.794/08) of the National Council for the Control of Animal Experimentation (CONCEA, Brazil). The protocol was approved by the Committee for Animal Use of the Universidade Federal do Rio de Janeiro (Permit Number: 161/18).

### Culture of Parasites

Infective promastigotes of *L. amazonensis* (MHOM/BR/75/Josefa) were obtained from infected BALB/c mouse lesions and were used until the 5^th^ culture passage as promastigotes at 26°C in M-199 medium (Cultilab) supplemented with 20% heat-inactivated fetal bovine serum (FBS) (Cultilab).

### *In vivo* Infection and Treatment

BALB/c mice were infected subcutaneously in the right hind footpad with 2×10^6^ stationary-phase promastigotes of *L. amazonensis* in 20 µl PBS. The following antibodies were administered intraperitoneally at 100 µg in 100 µl PBS; anti-PD-L1 (BMS-936559, Bristol-Myers Squibb), anti-PD-L2 (B7-DC, clone TY25, catalog # BE0112, Bioxcell), and anti-PD-1 (CD279, clone RMP1-14, catalog # BE0146, Bioxcell). The first injection was given at 7 days post-infection. Two treatment protocols were assessed: (i) inoculation once a week for 49 days with a total of 6 doses; and (ii) twice a week for 56 days with a total of 12 doses. Control animals received 100 μl PBS intraperitoneally also at 7 days post-infection and in accordance with the two treatment protocols. For both treatments, the last dose was administered 5 days prior to the euthanasia of the animals. Footpad thickness was measured weekly by using a direct-reading Vernier caliper.

### Parasite Load Quantification

After the mice were euthanized, the infected paws were removed, weighed, macerated with a tissue mixer, the homogenate diluted into 96-well culture plates (Jet Biofil, China) and incubated at 26°C for 7 and 15 days. Promastigote cultures were examined via optical microscope (Olympus, Japan), and the last well containing promastigotes in the limiting dilution assay was recorded to calculate the parasite load.

### Cell Staining for Flow Cytometry

Lymph nodes were removed and macerated with a tissue mixer; cells (1×10^6^/well in a 24-well plate) were cultured for 4 h at 37°C with Ionomycin (10 ng/ml, Sigma-Aldrich), Brefeldin (5 mg/ml, Biolegend) and PMA (phorbol 12-myristate 13-acetate, 10 ng/ml, Sigma-Aldrich). All centrifugation steps were performed at 4°C. Cells were washed with PBS and blocked with 50 µl/well (Human FcX, BioLegend) for 15 min. After which, 50 µl/well of the staining antibody pool was added and incubated for 30 min at 4°C. Cells were washed with buffer solution (PBS with 5% FBS) at 400 g for 5 min, then fixed and permeabilized (FoxP3 permeabilization/fixation kit; eBiosicence) according to the manufacturer’s protocol. Cells were washed again with buffer solution and resuspended in the same solution. For intracellular staining, the following antibodies were added (all used at 0.1 μg/ml) and incubated for 1 h at 4°C in the dark: CD3 (anti-CD3-APC-780; clone 145-2C11, Biolegend), CD4 (anti-CD4-PE-Cy7; clone RM4-5, Biolegend), CD8 (anti-CD8-PerCP; clone 53-6.7, Biolegend), PD-1 (anti-PD-1-FITC; clone J43, eBiosciences), CD25 (anti CD25-PE; clone P4A10, eBioscience), and intracellular IFN-γ (anti-IFN-γ-APC; clone XMG1.2, eBiosciences) and FoxP3 (clone FJK-16s; eBioscience). Cells were washed and resuspended in 150 μl buffer solution, and stored in the dark at 4°C until acquisition. Cells were analyzed on a BD FACS CANTO II flow cytometer; data from 100,000 events were captured from cells acquired on CD3^+^ and analyzed with FlowJo^®^ software (BD-Becton, Dickinson & Company).

### Analysis of Cytokines

The supernatants of the paw maceration were assayed for IL-4, IL-10, and TGF-β cytokines by specific ELISAs using a standard protocol (BD OptEIA).

### Dosage of Immunoglobulins

Soluble *L. amazonensis* antigen (LaAg) was obtained from stationary phase promastigotes, which were washed 3 times in PBS, freeze-thawed for 3 cycles, lyophilized, stored at −20°C and reconstituted with PBS just prior to use. The 96-well plates were coated with LaAg (1 μg/well) overnight at 4°C, blocked with PBS/5% milk/0.05% Tween 20 (Sigma-Aldrich) for 2 h, and washed 3 times with PBS/0.05% Tween 20. The mouse serum samples (1:250 diluted in PBS/5% milk/0.05% Tween 20) were added. The plates were incubated at room temperature for 1 h and washed with PBS/0.05% Tween 20. Anti-IgM or anti-IgG-HRP (Southern Biotech) were added (1:2000) for 1 h at room temperature. After washing, the color was developed with a TMB solution (Life Technologies) and stopped with 1 M HCl.

### Data Analysis

Results are expressed as mean ± SD with confidence level *p* ≤ 0.05. For lesion development analysis, a two-way ANOVA with a Bonferroni post-test was used. For multiple comparisons, a one-way ANOVA followed by Tukey pairing was performed. Paired t-test analysis was done as indicated in the figure legends. Data analysis was performed using GraphPad Prism^®^ 5.00 software.

## RESULTS

### *L. amazonensis* infection induces PD-1 expression on CD4^+^ and CD8^+^ T cells

Previous studies by our group have demonstrated that in footpads and lymph nodes of *L. amazonensis*-infected C57BL/6 mice, there is increased expression of PD-1 on CD4^+^ T cells in comparison to naïve uninfected mice.^21^ Given that the disease profile of *L. amazonensis* infection in C57BL/6 and BALB/c mice are different, in that BALB/c mice develop more progressive disease than C57BL/6 mice, we decided to evaluate the expression of PD-1 in lymph node cells of *L. amazonensis*-infected BALB/c mice. The data showed an increase in the percentage (Fig. 1a, c) and absolute number (Fig. 1b) of PD-1^+^ CD4^+^ and CD8^+^ T cells in infected compared to uninfected BALB/c mice.

**Fig. 1:**
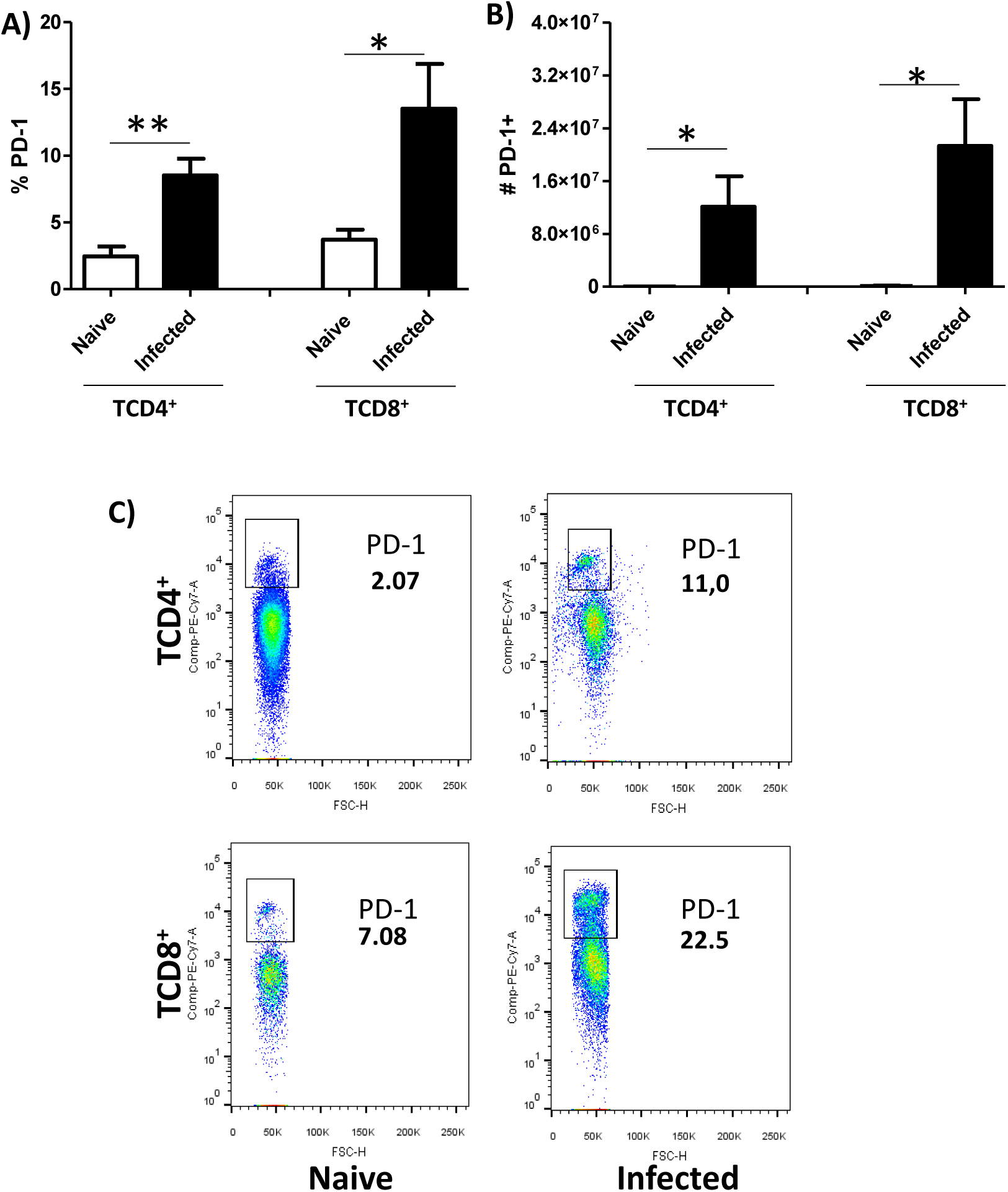
Expression of PD-1 on CD4^+^ and CD8^+^ T cells in *L. amazonensis*-infected BALB/c mice. Lymphocytes from the macerated draining popliteal lymph node of an *L. amazonensis*-infected paw were collected at 56 days post-infection. Uninfected mice were used as control. **a-b** Percentage and absolute number of PD-1^+^CD4^+^ T cells and PD-1^+^CD8^+^ T cells. **c** Dot plots showing PD-1 expression (PE-Cy7-PD-1^+^, FSC-cell volume). *p<0.05, **p<0.0375 (T Test). The data show means ± standard deviations; *n* = 8-13.

### Anti-PD-1 or anti-PD-L1 MoAb treatment reduces parasite loads without affecting lesion growth in mice

As we observed that PD-1 was upregulated during *L. amazonensis* infection, we next assessed the effect of treatment with the anti-PD-1, anti-PD-L1, and anti-PD-L2 blocking antibodies on the disease profile. First, *L. amazonensis-*infected BALB/c mice were treated with the individual MoAbs (100 μg each/mice) once weekly, beginning at 7 days post-infection, receiving a total of 6 doses. The footpad thickness was measured weekly, and the parasite load was determined after 49 days of treatment. We found that this therapy was not effective in modifying the lesion development profile (Suppl. Fig. 1a) or the parasite load (Suppl. Fig. 1b) compared to control mice injected with PBS.

In the second treatment protocol, mice were given individual MoAbs twice weekly, beginning at 7 days post-infection and receiving a total of 12 doses during the 56 day observation period. We found that although the lesion sizes (Fig. 2a) were increased during anti-PD1 and anti-PD-L1 treatment, the parasite loads were significantly decreased after anti-PD-1 or anti-PD-L1 treatment (Fig. 2b), Suppl. Fig. 2b). Anti-PD-L2 treatment did not show any significant effect (Suppl. Fig. 2). These results suggest the possibility of using the anti-PD-1 and anti-PD-L1 MoAb therapies to decrease parasitic load.

**Fig. 2:**
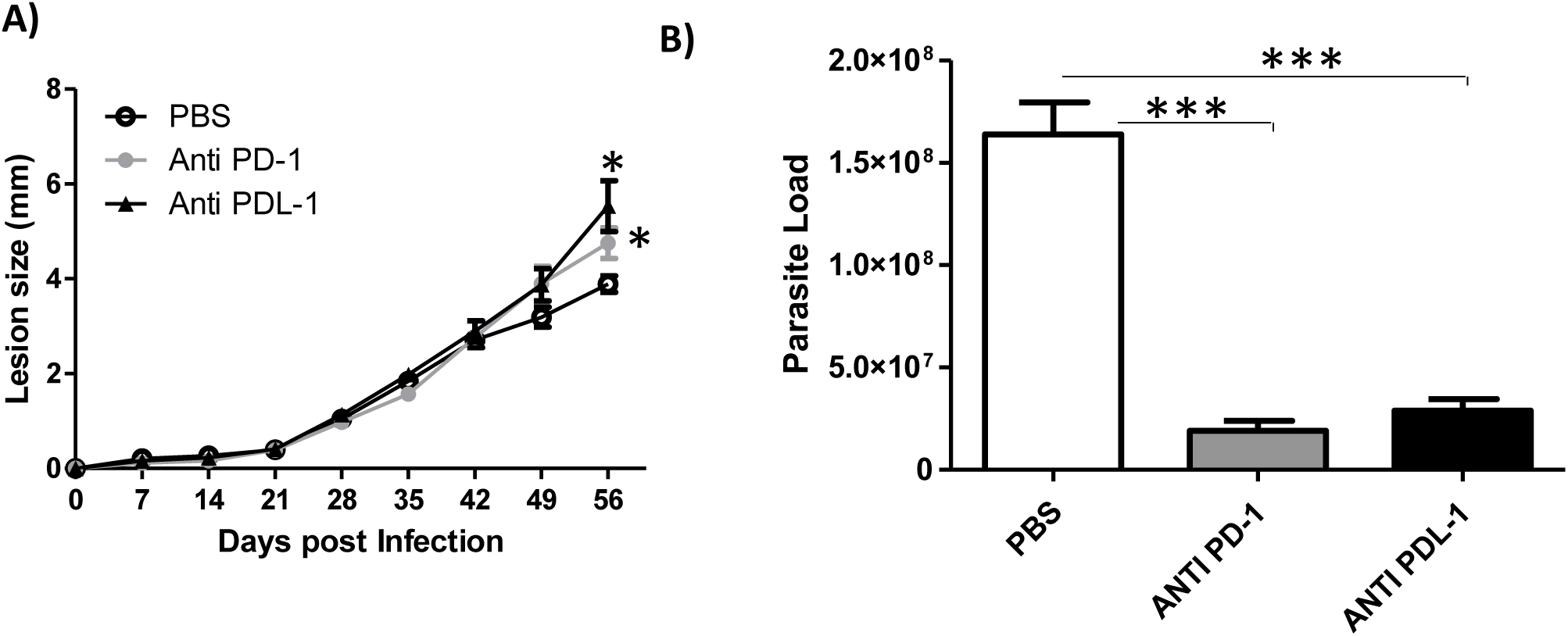
Lesion development and parasite load in *L. amazonensis*-infected BALB/c mice. Mice were infected *with L. amazonensis* promastigotes (2×10^6^) and treated with either anti-PD-1 or anti-PD-L1 MoAbs (100 μg/dose), administered twice a week intraperitoneally, beginning at 7 days post-infection, for 56 days. **a** Lesion size. **b** Parasite load. *p<0.05, ***p<0.0001 (t-test). The data (means ± standard deviations; *n* = 5) are representative of two independent experiments producing the same result profile.

### Induction of IFN-γ from CD4^+^ and CD8^+^ T cells after MoAb treatment

Focusing on the twice a week treatment protocol, we then examined the mechanism underlying parasite load reduction in anti-PD-1 or anti-PD-L1-treated mice by measuring the IFN-γ production of CD8^+^ and CD4^+^ T cells from the draining lymph nodes. It is documented in murine models of *L. major* infection that the expression of cytokines such as IL-12 and IFN-γ by Th1 contributes to host protection, whereas IL-4, IL-5, and IL-10 expression by Th2 contributes to host susceptibility.^22,23^ In the murine *L. amazonensis* infection model, we and others have shown that impaired IFN-γ production and insufficient macrophage activation favor parasite survival and persistence.^22-25^ In our treatment studies, we found more CD3^+^CD8^+^ T cells in the *L. amazonensis-*infected mice compared to uninfected mice (Fig. 1a), but there were no significant difference in the percentage and absolute number of CD3^+^CD8^+^ T cells between the MoAb-treated and the PBS-injected groups (Fig. 3a, b). However, both, percentage and absolute number of IFN-γ-producing CD3^+^CD8^+^ T cells (IFN-γ^+^CD8^+^ T cells) were significantly increased in anti-PD-1 or anti-PD-L1-treated mice when compared to the PBS-injected group (Fig. 3c, d, e).

**Fig. 3:**
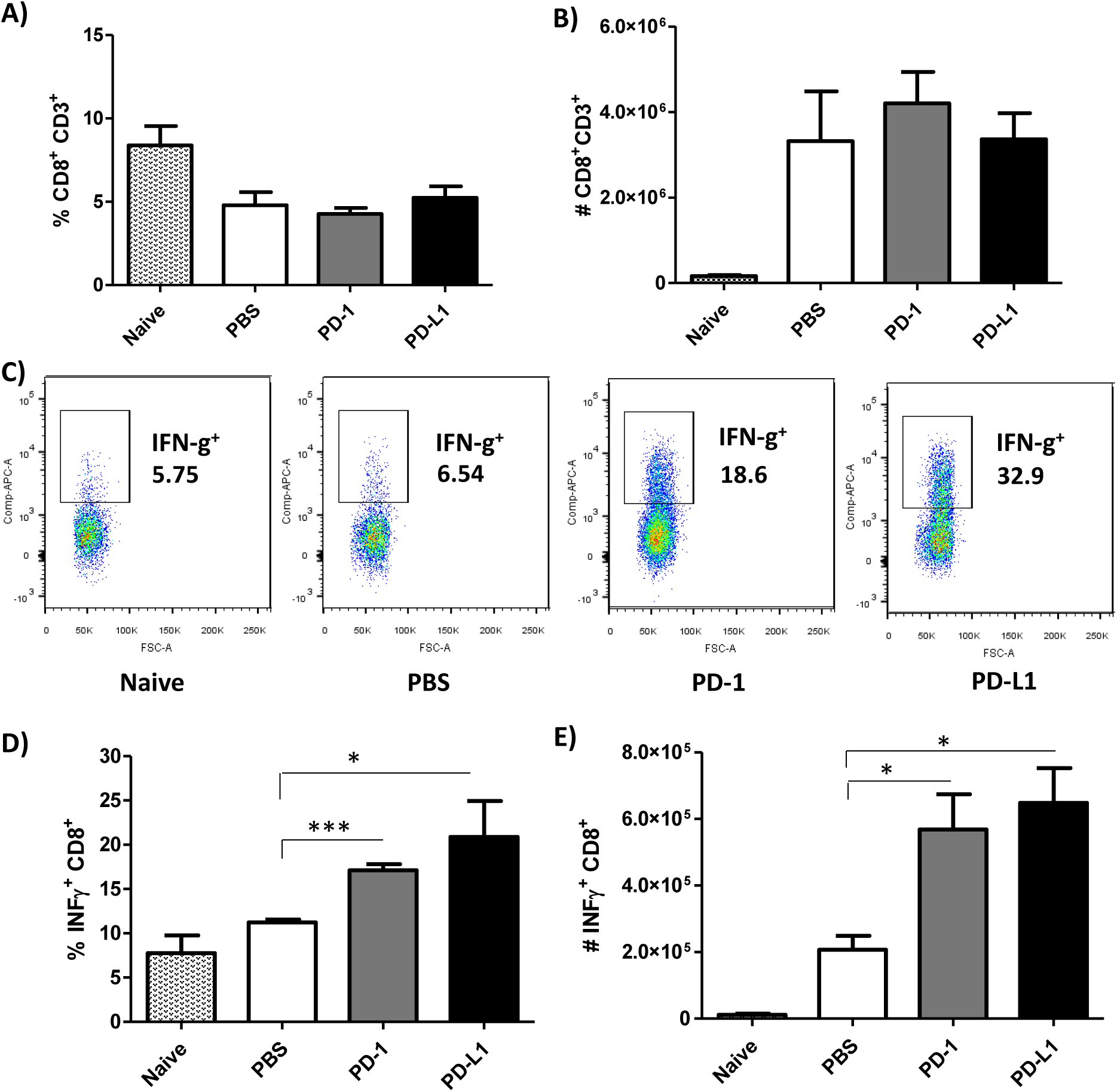
Effect of anti-PD-1 and anti-PD-L1 MoAbs on the percentage and number of IFN-γ^+^CD8^+^ T cells. Lymphocytes were collected from the draining lymph node of an *L. amazonensis*-infected paw after 56 days of treatment with anti-PD-1 or anti-PD-L1, administered twice a week intraperitoneally beginning 7 days after infection. **a** Percentage of CD3^+^CD8^+^ T cells. **b** Number of CD3^+^CD8^+^ T cells. **c** Dot plots showing IFN-γ expression (APC-IFN-γ, FSC-cell volume). **d** Percentage of IFN-γ^+^CD8^+^ T cells. **e** Number of IFN-γ^+^CD8^+^ T cells. *p<0.05, ***p<0.0001, (T Test (**d**), ANOVA (**e**)). Naive = mice without infection and therapy, PBS = infected mice injected with PBS on treatment days, PD-1 = mice infected and treated with anti-PD-1 (100 μg/dose), PD-L1= mice infected and treated with anti-PD-L1 (100 μg/dose). Data (means ± standard deviations; *n* = 5) are representative of three independent experiments producing the same result profile.

As the percentage and number of PD-1^+^CD8^+^ T cells were increased in *L. amazonensis* infection (Fig. 1), we next examined whether the IFN-γ production could also be affected in the CD8^+^ T cells lacking PD-1 (PD-1^−^CD8^+^) during MoAb treatment. Similar to the total CD3^+^CD8^+^ T cells, the PD-1^−^CD8^+^ T cells of both groups of MoAb-treated mice presented a higher percentage of IFN-γ expression, but there was not a significantly higher number of IFN-γ^+^PD-1^−^CD8^+^ T cells compared to the PBS-injected group (Fig. 4a, b, c).

**Fig. 4:**
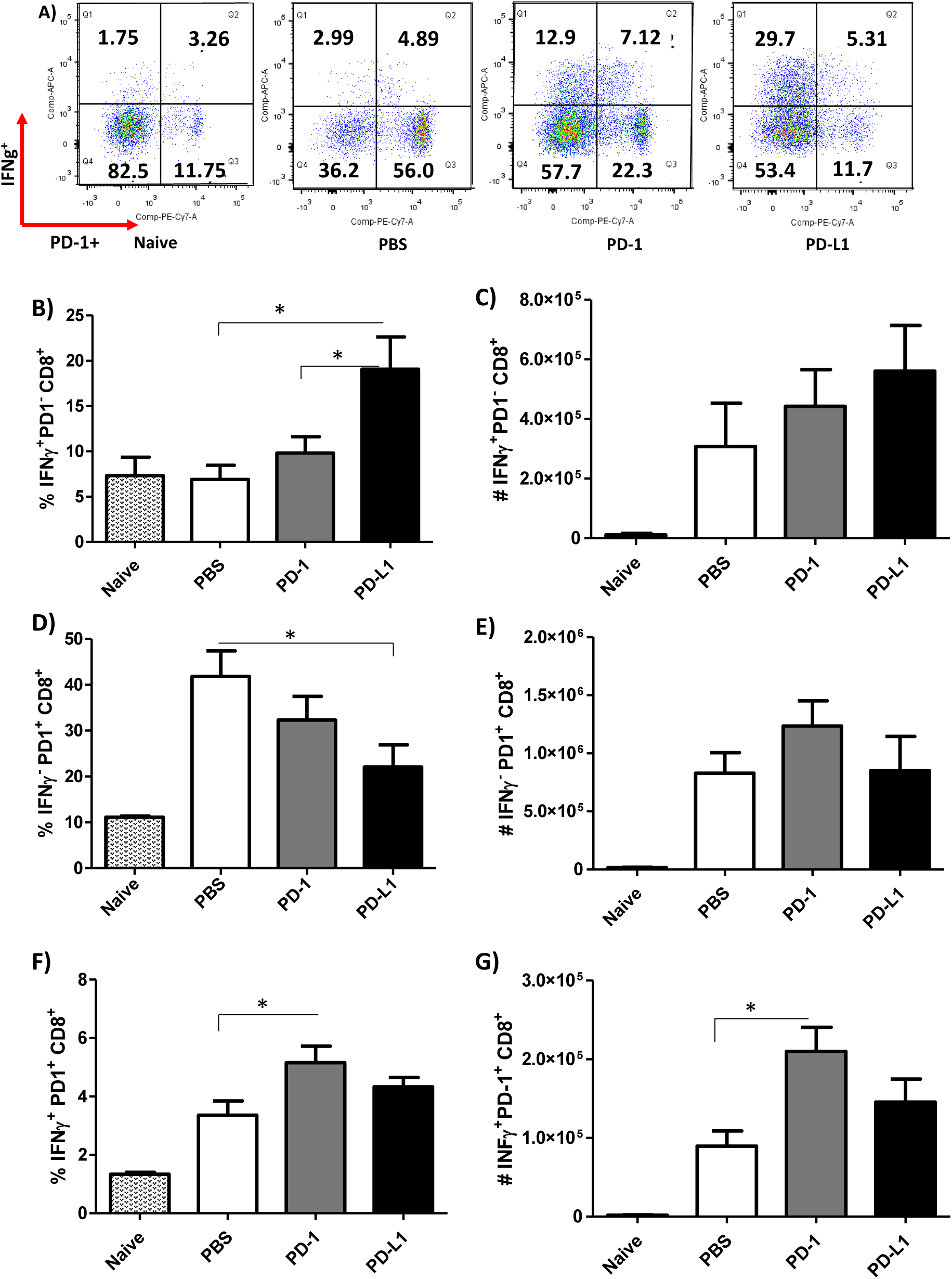
Increase of IFN-γ^+^CD8^+^ T cells after anti-PD-1 and anti-PD-L1 MoAb treatment. Lymphocytes were collected from the popliteal lymph node of an *L. amazonensis*-infected paw after approximately 2 months of treatment with anti-PD-1 or anti-PD-L1, administered twice a week intraperitoneally beginning 7 days after infection. **a** Dot plot IFN-γ and PD-1 expression (APC-IFN-γ, PE-Cy7-PD-1). **b** Percentage of IFN-γ^+^PD-1^−^CD8^+^ T cells. **c** Number of IFN-γ^+^PD-1^−^ CD8^+^T cells. **d** Percentage of IFN-γ^−^PD-1^+^CD8^+^T cells. **e** Number of IFN-γ^−^PD-1^+^ CD8^+^T cells. **f** Percentage of IFN-γ^+^PD-1^+^CD8^+^ T cells. **g** Number of IFN-γ^+^PD-1^+^CD8^+^ T cells. *p <0.05, (T Test). Naive= mice without infection and therapy, PBS = infected mice injected with PBS on treatment days, PD-1 = mice infected and treated with anti-PD-1 (100 μg/dose), PD-L1= mice infected and treated with anti-PD-L1 (100 μg/dose). The data (means ± standard deviations; *n*= 5) are representative of three independent experiments producing the same result profile.

Regarding PD-1^+^CD8^+^ T cells not expressing IFN-γ (IFN-γ ^−^PD-1^+^CD8^+^), there was a significant decrease in the percentage of these cells in anti-PD-L1-treated mice compared to the PBS-injected group, without any effect on the absolute numbers (Figure 4 d, e). We did observe an increase in the IFN-γ-expressing PD-1^+^CD8^+^ T cells in both the percentage and absolute number of anti-PD-1-treated mice, but there was no significant difference in anti-PD-L1-treated mice compared to the PBS-injected group (Fig. 4f, g).

We also investigated the profile of CD4^+^ T cells and found no major differences between the MoAb-treated and PBS-injected groups regarding the percentage and number of CD3^+^CD4^+^ T cells (Fig. 5a, b). Like the CD3^+^CD8^+^ T cells, the percentage of CD4^+^ T cells producing IFN-γ was significantly increased after treatment with anti-PD-1 and anti-PD-L1 (Fig. 5c, d), however, a significant increase in the number of IFN-γ^+^-producing CD4^+^ T cells was observed in the anti-PD-1 treatment (Fig. 5e).

**Fig. 5:**
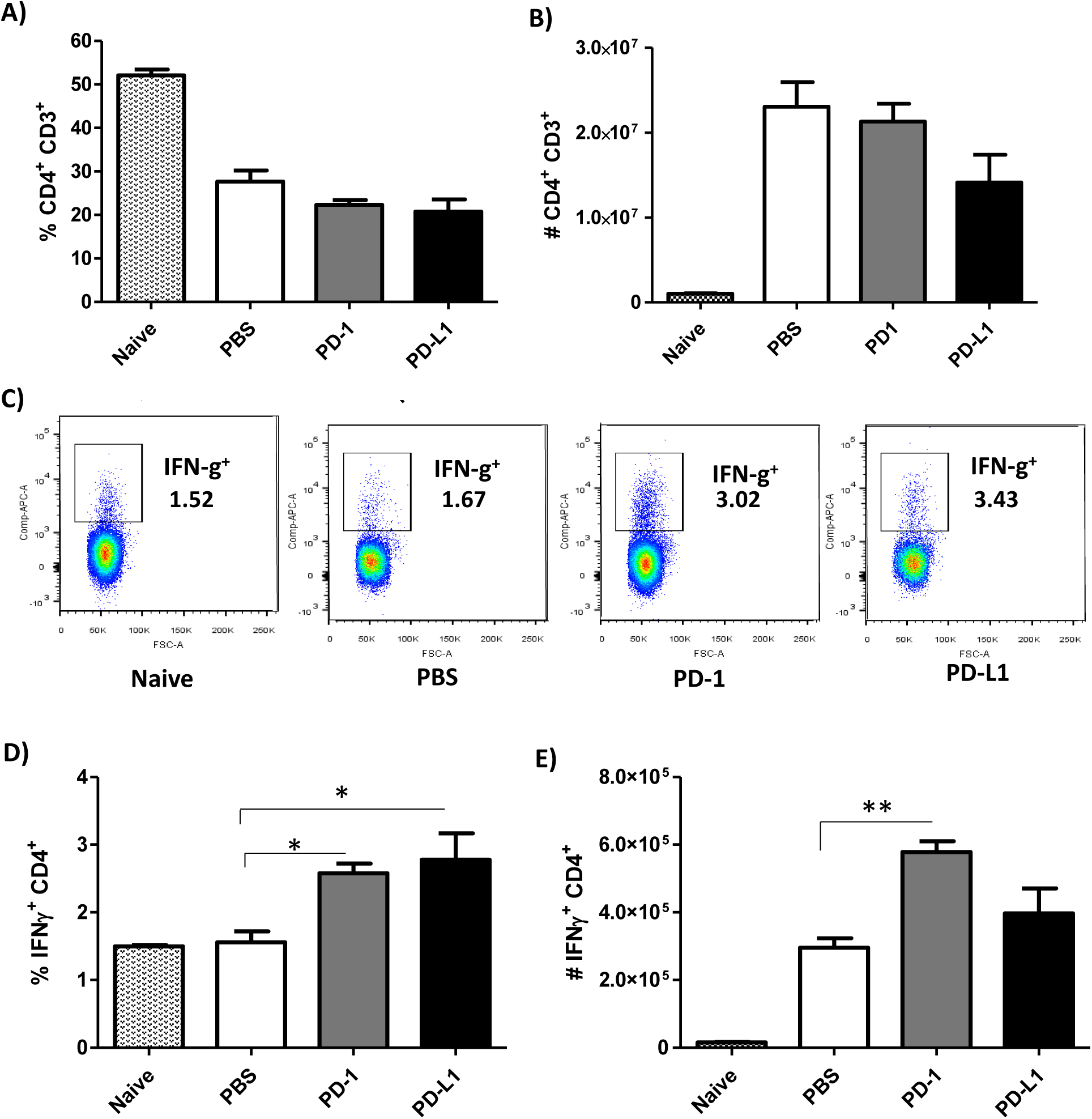
Increase of IFN-γ^+^CD4^+^ T cells after anti-PD-1 and anti-PD-L1 MoAb treatment. Lymphocytes were collected from the popliteal lymph node of an *L. amazonensis*-infected paw after approximately 2 months of treatment with anti-PD-1 or anti-PD-L1, administered twice a week intraperitoneally beginning after 7 days of infection. **a** Percentage of CD3^+^CD4^+^ T cells. **b** Number of CD3^+^CD4^+^ T cells. **c** Dot plot of IFN-γ expression (APC-IFN-γ, FSC-cell volume). **d** Percentage of IFN-γ^+^CD4^+^ T cells. (E) Number of IFN-γ^+^ CD4^+^ T cells. *p<0.05, **p<0.0375 (T Test (**d**) and ANOVA (**e**)). Naive = mice without infection and therapy, PBS = infected mice injected with PBS on treatment days, PD-1 = mice infected and treated with anti-PD-1 (100 μg/dose), PD-L1 = mice infected and treated with anti-PD-L1 (100 μg/dose). The data (means ± standard deviations; *n* = 5) are representative of three independent experiments producing the same result profile.

Unlike the PD-1^−^CD8^+^ T cells, MoAb treatment had no effect on the IFN-γ production by PD-1^−^CD4^+^ T cells (Fig. 6a, b, c). Again, opposite to what was observed in the IFN-γ^−^PD-1^+^CD8^+^ T cells, there was a significantly higher percentage of IFN-γ^−^ PD-1^+^CD4^+^ T cells in anti-PD-L1-treated mice, with no difference on the absolute number of these cells (Fig. 6d, e). Additionally, a significant increase in the percentage of IFN-γ-producing PD-1^+^CD4^+^ T cells (IFN-γ^+^PD-1^+^CD4^+^) was found for both MoAb-treated groups compared to the PBS-infected group, which was not observed in the number of cells (Fig. 6f, g).

**Fig. 6:**
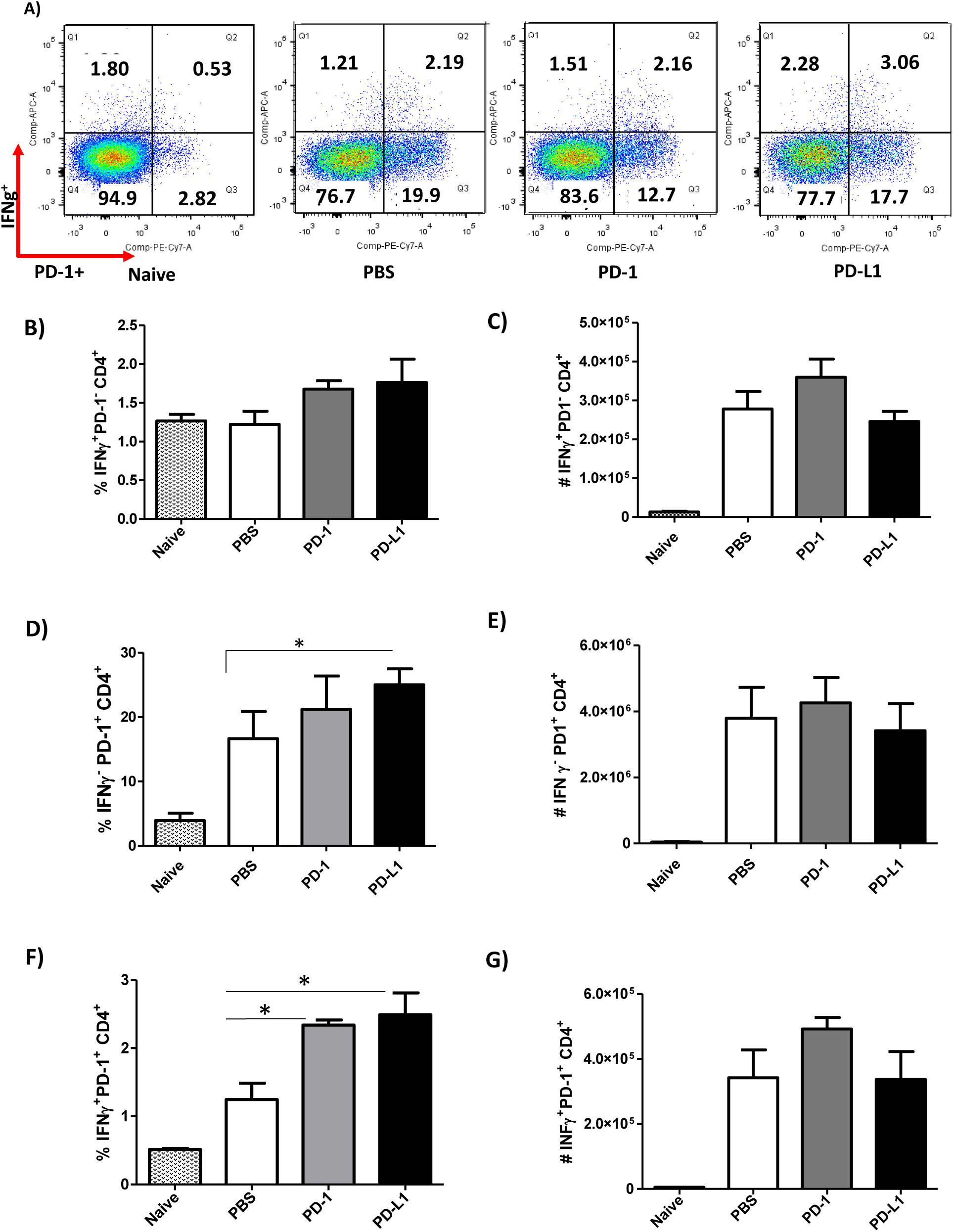
Increase in the percentage of IFN-γ^+^PD-1^+^CD4^+^ T cells after anti-PD-1 and anti-PD-L1 MoAb treatment. Lymphocytes were collected from the popliteal lymph node of an *L. amazonensis*-infected paw after approximately 2 months of treatment with anti-PD-1 or anti-PD-L1, administered twice a week intraperitoneally beginning after 7 days of infection. **a** Dot plot of IFN-γ and PD-1 expression (APC-IFN-γ, PE-Cy7-PD-1). **b** Percentage of IFN-γ^+^PD-1^−^CD4^+^ T cells. **c** Number of IFN-γ^+^PD-1^−^CD4^+^ T cells. **d** Percentage of IFN-γ^−^PD-1^+^ CD4^+^ T cells. **e** Number of IFN-γ^−^PD-1^+^CD4^+^ T cells. **f** Percentage of IFN-γ^+^PD-1^+^CD4^+^ T cells. **g** Number of IFN-γ^+^PD-1^+^ CD4^+^ T cells. *p <0.05 (T Test (**d**) and ANOVA (**f**)). Naive = mice without infection and therapy, PBS = infected mice injected with PBS on treatment days, PD-1= mice infected and treated with anti-PD-1 (100 μg/dose), PD-L1= mice infected and treated with anti-PD-L1 (100 μg/dose). The data (means ± standard deviations; *n* = 5) are representative of three independent experiments producing the same result profile.

Altogether, our results suggest that anti-PD-1 and anti-PD-L1 MoAb treatment stimulated the production of IFN-γ in both CD4^+^ and CD8^+^ T cells, which may be one of the possible mechanisms for the control of parasite load.

### Anti-PD-1 treatment reduces IL-4 and TGF-β

As IFN-γ production was induced by the MoAb treatment, we tested the effect of treatment on the modulation of the cytokines, IL-4, IL-10 and TGF-β, at the site of *L. amazonensis* infection. Only anti-PD-1 treatment significantly decreased IL-4 (Fig. 7a) and TGF-β (Fig. 7b) production *in situ* compared to the PBS-injected control group. No alteration in the IL-10 levels was found after MoAb treatment (Fig. 7c). Finally, we demonstrated that treatment with the MoAbs did not affect the production of anti-*Leishmania* specific IgM or IgG antibodies as assessed in the blood sera (Suppl. Fig. 4a, b). Altogether, our results suggest that anti-PD-1 MoAb treatment modulates the production of IFN-γ in CD4^+^ T and CD8^+^ T cells, but anti-PD-L1 only affects CD8^+^ T cells.

**Fig. 7:**
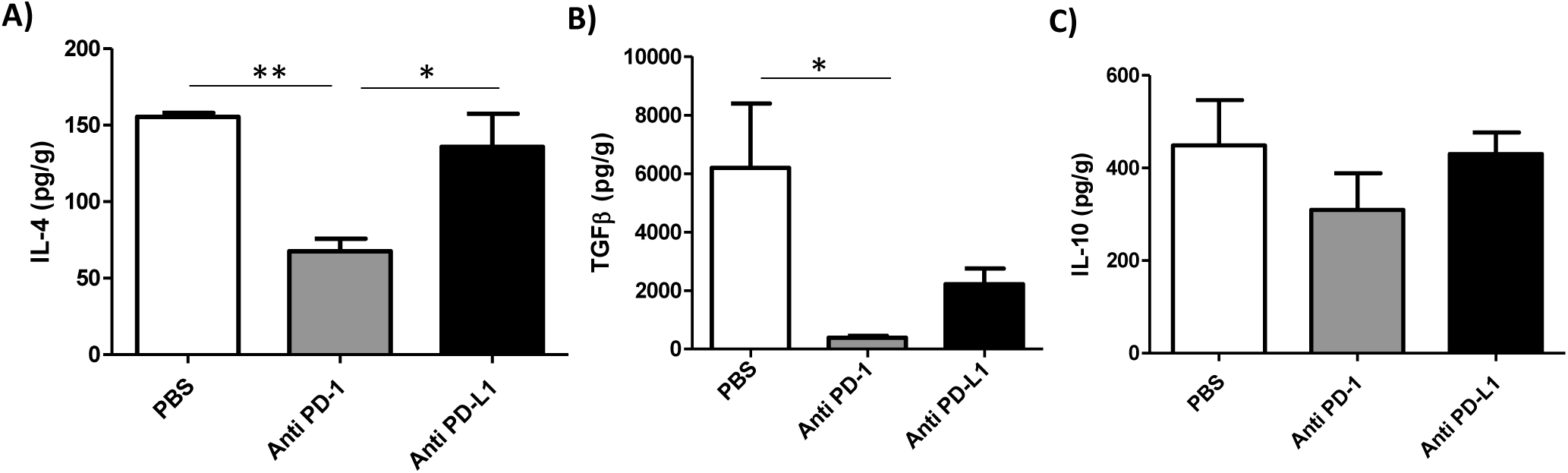
Selective reduction of IL-4 and TGF-β in MoAb-treated groups. *L. amazonensis*-infected paws were collected and macerated after approximately 2 months of treatment with anti-PD-1 or anti-PD-L1, administered intraperitoneally twice a week beginning after 7 days of infection, and the supernatant was analyzed by ELISA: **a** IL-4, **b** TGF-β, **c** IL-10. *p<0.05, **p<0.0375 (ANOVA). The data (means ± standard deviations; *n* = 5) are representative of two independent experiments producing the same result profile.

## DISCUSSION

The use of leukocyte receptor blockers in immunotherapy has been extensively studied in oncology.^26^ One of the clinical trials concerning the systemic administration of therapeutic antibodies to block PD-1 or PD-L1 has produced promising results for the treatment of several tumors.^27,28^ However, the use of this therapy is still relatively limited in non-healing leishmaniasis.

*L. amazonensis* infection in C57BL/6 mice induced expression of PD-1 on CD4^+^ T cells (de Matos Guedes et al, submitted); however, in BALB/c mice the expression of PD-1 was induced on both CD4^+^ and CD8^+^ T cells prompting a more severe impairment of IFN-γ production. For *L. major* infection in arginase-deficient mice, deficiency in T cell activation resulted in increased PD-1 expression, impairing the immune response and inducing T cell exhaustion.^29^ In dogs injected with *L. infantum* antigens, PD-L1/PD-1 blockade with specific antibodies recovered the proliferation of CD4^+^ and CD8^+^ T cells, in addition to the production of IFN-γ by CD4^+^ T cells.^30^ In studies of *L. donovani* infection in BALB/c mice, the parasite induced the initial expansion of IFN-γ-producing CD4^+^ and CD8^+^ T cells in the acute phase of the disease, the frequency of which was reduced after 21 days post-infection even with a robust parasite presence. In this model, blocking PD-L1 resulted in the restoration of CD4^+^ and CD8^+^ T cell responses, leading to a reduction in parasite load.^31^ In view of these studies, our results presented herein show that *L. amazonensis* infection interferes in the production of IFN-γ by both CD4^+^ and CD8^+^ T cells.

Recently, the presence of PD-1 and PD-L1 was detected in a patient with diffuse cutaneous leishmaniasis caused by *L. amazonensis.*^32^ Based on the induction of PD-1 and PD-L1 by *Leishmania* infection, the blocking of these molecules may be a new strategy to treat leishmaniasis.

The use of anti-PD-1 and anti-PD-L1 antibodies in clinical cancer treatment studies, such as in pancreatic tumor, were administered at doses between 10 and 200 μg every three days.^33-35^ In our study, we tested a lower therapeutic dose of 100 µg of the anti-PD-L1, anti-PD-L2 and anti-PD-1 antibodies once a week. However, our data revealed that this was insufficient in reducing the lesion size or parasite load. Therefore, the dosage of the treatment, in terms of the concentration and frequency of administration are important considerations. Thus, we increased the administration to twice a week, and observed that although the lesion size was increased, the parasite load in the infected footpads, in the spleen and in the draining lymph nodes that received therapy with anti-PD-1 and anti-PD-L1 were controlled.

In *L. major* infection, only treatment with 1 mg/dose of the anti-PD-1 antibody weekly in infected arginase-deficient mice led to complete resolution of the chronic skin lesion and resistance to infection.^29^ A similar result was found when using the anti-PD-L1 antibody in mice challenged with *L. donovani* amastigotes, as these mice showed a reduction of up to 87% of the parasite load in the spleen.^36^ The susceptibility of the C57BL/10 and C57BL/6 mice to infection by *L. amazonensis* is related to the absence of Th1 type cellular immune response and not controlled exclusively by Th2 cells.^37,38^ In BALB/c mice, this susceptibility is related to Th2 type cellular immune response.^39,40^

In this study, we observed that the production of IFN-γ was higher in CD8^+^ T lymphocytes, both in percentage and number of cells after treatment with the anti-PD-1 and anti-PD-L1 MoAbs. We also detected an increase in the percentage of IFN-γ-producing CD4^+^ T lymphocytes after both treatments. However, when looking at the number of IFN-γ-producing CD4^+^ T cells only in mice treated with anti-PD-1 a significant increase was observed. These results suggest that treatment with anti-PD-1, acting directly on the lymphocytes, is more competent in invigorating CD4^+^ T lymphocytes than the anti-PD-L1 therapy, the target of which is in the antigen-presenting cells. Our data indicate that the increase of IFN-γ is one of the possible mechanisms of the therapeutic efficacy of these monoclonal antibody therapies, mainly by CD8^+^ T and partially by CD4^+^ T lymphocytes.

The PD-1/PD-L1 ratio is very important in suppressing the CD8^+^ T cell response during *L. donovani* infection.^36^ In a study of dogs with symptomatic visceral leishmaniasis by *L. infantum*, these animals were shown to have a five-fold reduction in the proliferative capacity of CD8^+^ T cells and a reduction of up to three-fold in the ability of these cells to produce IFN-γ. After administration of specific monoclonal antibody therapy, PD-1 blockade significantly increased the proliferative capacity of CD4^+^ and CD8^+^ T cell populations, and recovered IFN-γ production in the CD4^+^ population. In addition, T cell depletion during visceral leishmaniasis was associated with elevated expression of PD-1, which could be identified before the onset of the disease and is considered a determining factor for symptomatic onset.^30^ In another study of canine visceral leishmaniasis, it was observed that as the disease progressed, there was a decrease in CD4^+^ T cell proliferation and also a reduction of IFN-γ production in response to *L. infantum* antigens.^39^

These findings are reinforced by studies in which anti-PD-1 and anti-PD-L1 therapy reverts the ability of CD8^+^ T lymphocytes to produce IFN-γ as in the treatment of thyroid cancer,^40^ in chronic hepatitis B,^41^ and in HIV infection.^42^ Further studies should be performed to confirm whether T cell exhaustion occurs in *L. amazonensis* infection. However, other studies that have assessed how chronic infections can induce exhaustion, in addition to our results reporting the increased capacity of IFN-γ production after MoAbs therapy, this suggests that T cell exhaustion occurs in *L. amazonensis* infection.

Interestingly, our results indicate the MoAb potential to modulate the cytokines present at the lesion site in the mice, as the production of IL-4 and TGF-β were reduced in the group treated with the anti-PD-1 MoAb. The group treated with anti-PD-L1 MoAb showed a tendency in the reduction of TGF-β. It is already known that patients who progress to the disease have an increase in the production of IL-4 directed by a Th2 response and suppressor responses by the increase of TGF-β (*L. chagasi*) and IL-10 (*L. donovani*).^43,44^ In another study, TGF-β was able to upregulate the PD-L1 expression in dendritic cells, leading to T-cell anergy and diminished anti-tumor response.^45^ In visceral leishmaniasis, it has been observed that patients who had progression of the disease showed increased IL-10 production,^39^ which was not seen in our results, since the concentration of IL-10 showed a slight reduction in the MoAb-treated groups relative to the PBS control. Thus, our data support the hypothesis that IFN-γ produced by CD4^+^ T cells inhibits the development of a Th2 response by diminishing IL-4 and reducing TGF-β production. Moreover, it is important to point out that TGF-β is associated with increased parasite load^46^ and its reduction is directly related to parasite control. Hence, in our results, parasitic control may also be related to the decrease of TGF-β and PD-L1 expression.

We also evaluated if the humoral response was altered during MoAb treatment. Kima *et al.* showed that antibodies play a critical role in the pathogenesis and in the development of more significant lesions due to *L. amazonensis* infection, since the maintenance of infection by these parasites was impaired by the absence of circulating antibodies in the BALB/c model.^47^ Recently, we demonstrated that *L. amazonensis* infection in XID mice displayed smaller lesions and a decrease in IL-10 and total antibodies in comparison to WT mice suggesting the pathogenic role of B cells in *L. amazonensis* infection.^48^ Here, we demonstrated that treatment with both MoAbs did not affect the anti-*Leishmania* IgM and IgG antibodies levels.

In summary, our study suggests a potential use of monoclonal antibodies against PD-1 and PD-L1 in the treatment of cutaneous leishmaniasis caused by *L. amazonensis*. Our model, which employed a low dose treatment of anti-PD-1 or anti-PD-L1 showed therapeutic efficacy to control the parasite load in infected mice. We have also shown that this control is related to CD8^+^ T lymphocytes and, partially, to CD4^+^ T lymphocytes, producing IFN-γ. These findings could potentiate a combined therapy using anti-PD-1 or anti-PD-L1 antibodies and the current standard therapies against leishmaniasis, which could be particularly important for diffuse cutaneous leishmaniasis treatment, a disease that is refractory to conventional treatment.

## ACKNOWLEDGEMENTS

We would like to thank Bristol-Myers Squibb for the donation of anti-PD-L1, through the sharing of these technologies we can hopefully present a new cure for leishmaniasis. This work was supported by Fundação Carlos Chagas Filho de Amparo à Pesquisa do Estado do Rio de Janeiro (FAPERJ), Conselho Nacional de Desenvolvimento Científico e Tecnológico (CNPq), and Coordenação de Aperfeiçoamento de Pessoal de Nível Superior (CAPES), Finance Code 001.

## SUPPLEMENTAL FIGURES

**Suppl. Fig. 1:**
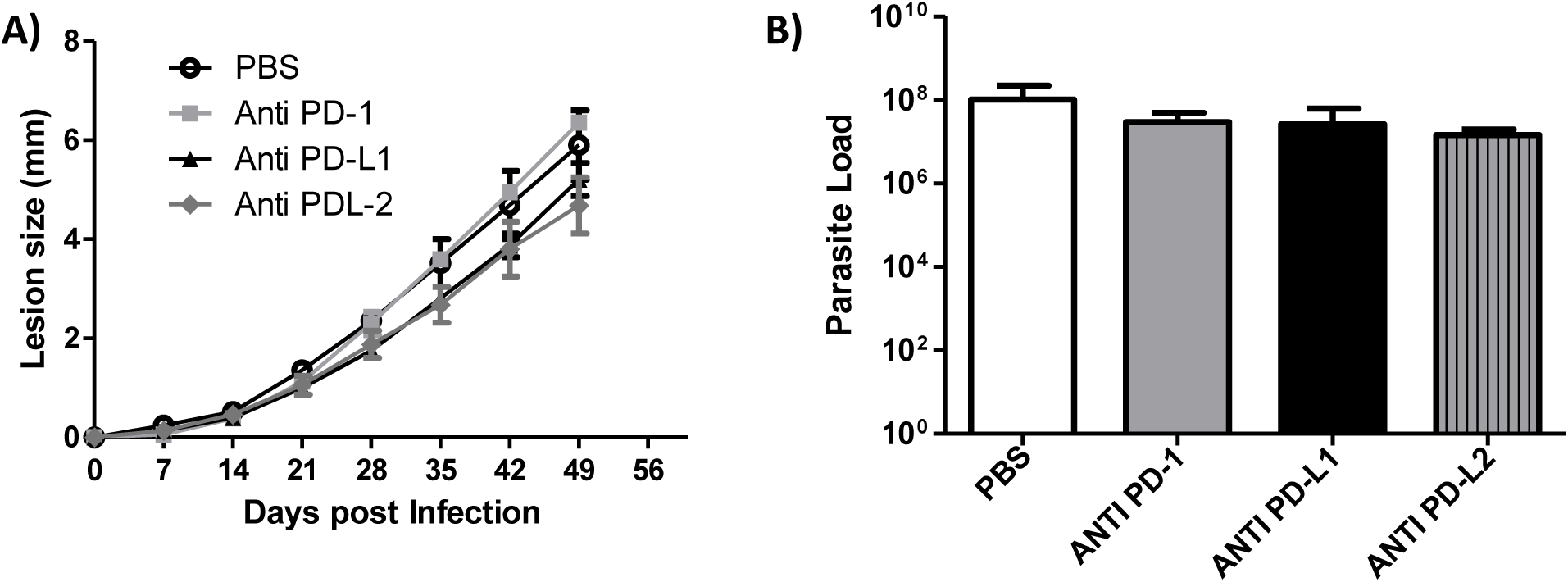
Ineffectiveness of treatment with a weekly dose of specific antibodies. Mice were infected in the footpad with *L. amazonensis* promastigotes (2×10^6^) and treated with anti-PD-1, anti-PD-L1 or anti-PD-L2, all at a 100 μg/dose, administered once a week intraperitoneally beginning at 7 days post-infection. (A) Progression of the lesion. (B) Parasite load. The data (means ± standard deviations; *n* = 5) are representative of an experiments.

**Suppl. Fig. 2:**
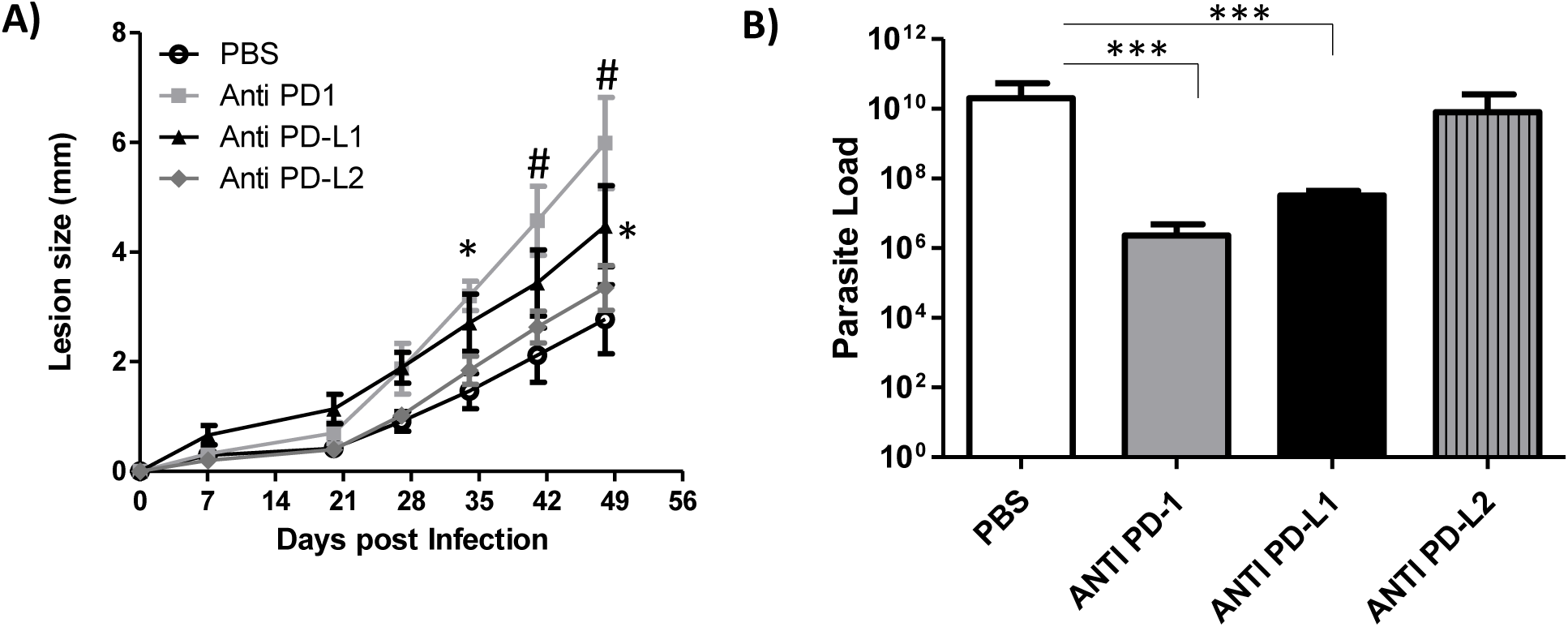
Unaltered lesion growth and control of parasite load from two weekly doses of specific antibodies. Mice were infected in the footpad with *L. amazonensis* promastigotes (2×10^6^) and treated with anti-PD-1, anti-PD-L1 or anti-PD-L2, all at a 100 μg/dose, administered twice per week intraperitoneally beginning at 7 days post-infection. (A) Progression of the lesion. (B) Parasite load of paw analyzed by limiting dilution. ***p <0.0001 (T Test). The data (means ± standard deviations; *n* = 5) are representative of an experiment.

**Suppl. Fig. 3:**
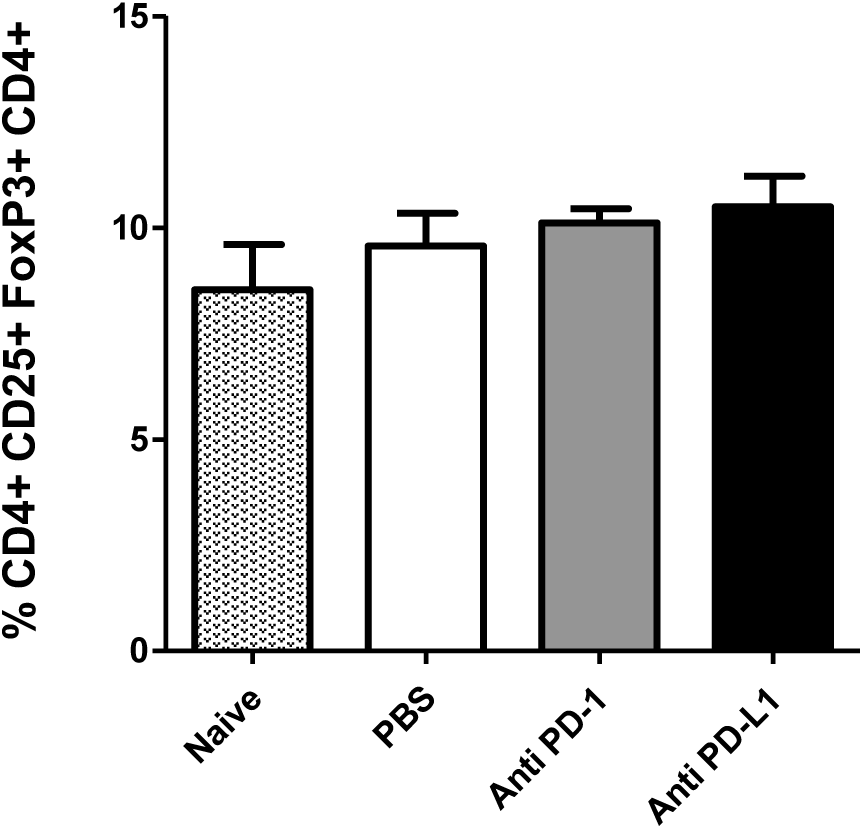
Treg cells in *L. amazonensis-*infected BALB/c mice. Mice were infected in the footpad with *L. amazonensis* promastigotes (2×10^6^) and treated with anti-PD-1, anti-PD-L1 or anti-PD-L2, all at a 100 μg/dose administered twice per week intraperitoneally beginning at 7 days post-infection. Naive mice were used as control. Percentage of Treg cells. The data (means ± standard deviations; *n* = 5) are representative of two independent experiments producing the same result profile.

**Suppl. Fig. 4:**
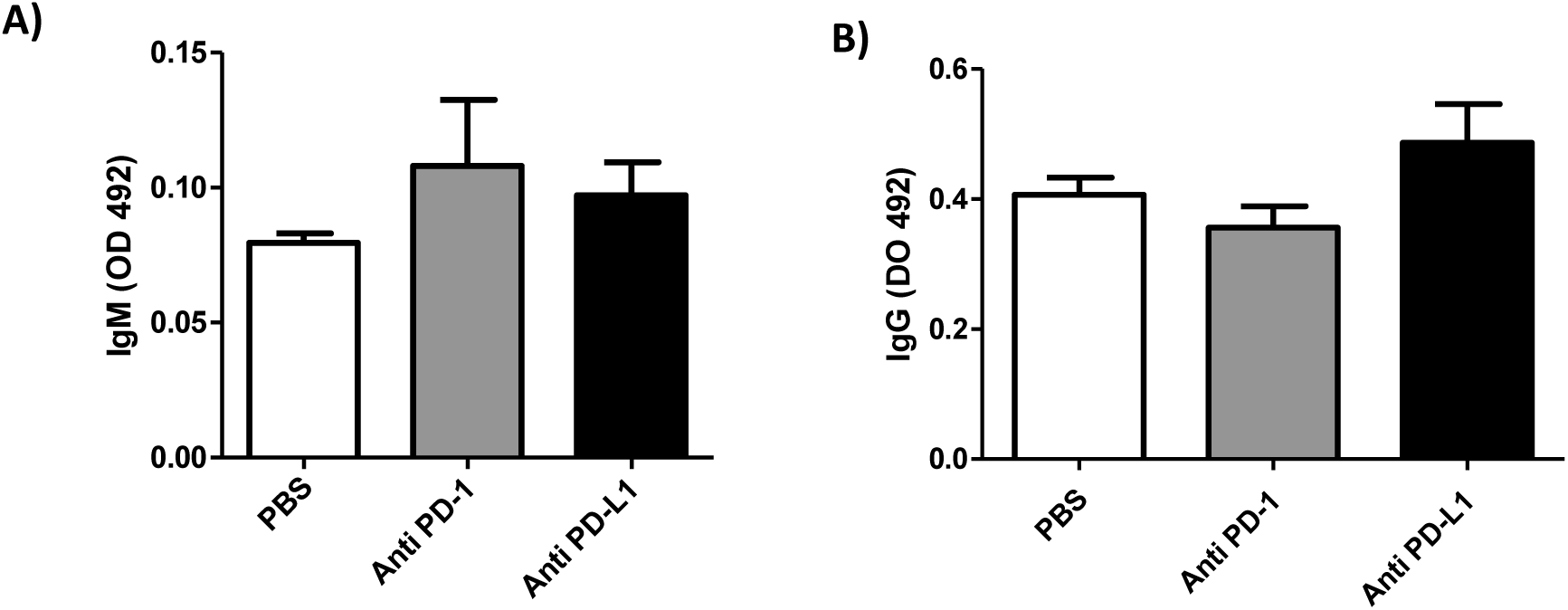
Effects of MoAb treatment on the specific anti-*Leishmania* immunoglobulins. (A) Specific IgM and (B) IgG antibodies were detected in the serum (diluted 1:500) of *L. amazonensis*-infected mice through ELISA using total *L. amazonensis* antigens (1 μg/well). Animals were treated for 49 days with antibodies administered twice a week intraperitoneally, starting at 7 days post-infection. Data (means ± standard deviations; *n* = 5) are representative of two independent experiments producing the same result profile.

